# Transcriptional characterization of the glial response due to chronic neural implantation of flexible microprobes

**DOI:** 10.1101/2021.05.31.446394

**Authors:** Kevin Joseph, Matthias Kirsch, Midori Johnston, Christian Münkel, Thomas Stieglitz, Carola A. Haas, Ulrich G. Hofmann

## Abstract

Long term implantation of (micro-)probes into neural tissue cause unique and disruptive responses to these foreign bodies. In this study, we present the transcriptional trajectory of glial cells responding to chronic implantation of flexible micro-probes for up to 18 weeks. Transcriptome analysis shows a rapid activation of microglial cells and a strong upregulation of reactive astrocytic genes, which is lost over the full duration of the implant period. Most interestingly, animals that were implanted for 18 weeks show a transcriptional profile similar to non-implanted controls, with increased expression of genes associated with wound healing and angiogenesis, which raises hope of a normalization of the neuropil to the pre-injury state when using flexible probes. Nevertheless, our data show, that a subset of genes upregulated after 18 weeks belong to the family of immediate early genes, which would indicate that structural and functional remodeling has not been completed at this time point. Our results confirm and extend previous work on the molecular changes resulting from the presence of intraneural probes and provide a rational basis for developing intervention strategies to control them.

## Introduction

Neural interfaces intended to interact with the central nervous system (CNS) are well established within fundamental neuroscience and aim at clinical applications (1–4). Accurate sampling of the electrophysiological activity of the local neuronal population is achieved by implanting penetrating probes into the brain parenchyma, sometimes equipped with a multitude of recording sites (5–7). Implantable neural interfaces can actively sample neuronal activity over extended periods of time and can, for instance, be used to study the neural circuitry involved in memory and learning, or the disruption of such pathways as a consequence of neurological disorders (8). Even though these neural interfaces are extremely promising, they have been long plagued by the loss of signal fidelity over chronic implantation time scales (9). To elucidate the mechanisms behind this loss, histological responses of the CNS to invasive implantation procedures associated with brain machine interfaces have been described by various authors over the years. The foreign body response of the brain upon implantation leads to a spatially confined inflammatory response and glial scarring, potentially leading to probe failure (10,11). It is therefore desirable to optimize probe designs towards attenuating the foreign body response in order to maintain high signal fidelity over chronic timescales (12,13). There has been a growing interest in the usage of flexible microelectrodes to minimize the glial activation around the recording sites to enable high quality recordings, which have proven to be promising as compared to rigid microelectrodes (14,15). However, there exists sparse information regarding the transcriptomic alterations due to the implantation of these flexible electrodes apart from a few studies (16,17), which would aid in identifying relevant genes and pathways amenable for targeting, by pharmacotherapeutics that can modulate pathways involved in the reactive response of the brain to a foreign body (18).

In this study, we extend the analysis of the effects of flexible probe implantation into the rodent neocortex over a chronic timescale of 18 weeks post implantation by using whole transcriptome analysis with the aim to identify crucial mechanisms which are involved in the cortical response to a foreign body. We report a significant increase of immune system regulation, extracellular matrix degradation and microglia/macrophage activation within 4 hours of implantation, coupled with the presence of reactive astrocytes. Following implantation and lasting 2 weeks, we see an upregulation of genes related to *Hmox1*^+^ macrophages, which has been previously reported to be responsible for inducing scar generating reactive astrocytes, mediated by Interleukin 10 signaling (19). This reactive astrocyte transformation gradually returns to the baseline homeostatic state over the implantation period, with no differences to control conditions within 18 weeks of implantation. Taken together, the microglia-astrocyte interaction seems to be an important point of intervention to modulate the formation of the glial scar, which would be the long-term enabler of improved brain machine interfaces.

## Results

### Acute injury initiates inflammatory microglial activation

To interrogate changes in gene expression due to acute injury as a result of the implantation trauma (**Fig. 1a**: Workflow), microarray-based transcriptome analysis was performed. Gene expression analysis revealed a clear difference between paired samples, with a clear distinction between the control and implanted samples. As one would expect to see from neural damage, a significant enrichment of matrix metalloproteinase genes (*Mmp12*, p_adj_<0.001; *Mmp14*,p_adj_=0.003) and genesets related to damage to the extracellular matrix (GO:0043312, NES=2.04, p_adj_=0.001; GO:0030198, NES=2.26, p_adj_<0.0001) was observed (**Fig. 1b-c)**. S*pp1,* also known as osteopontin (*Opn*), a pro inflammatory cytokine that plays an important role in tissue repair and extracellular matrix remodeling after injury was significantly up-regulated (p_adj_<0.0001), (41–44)).

**Figure 1:**
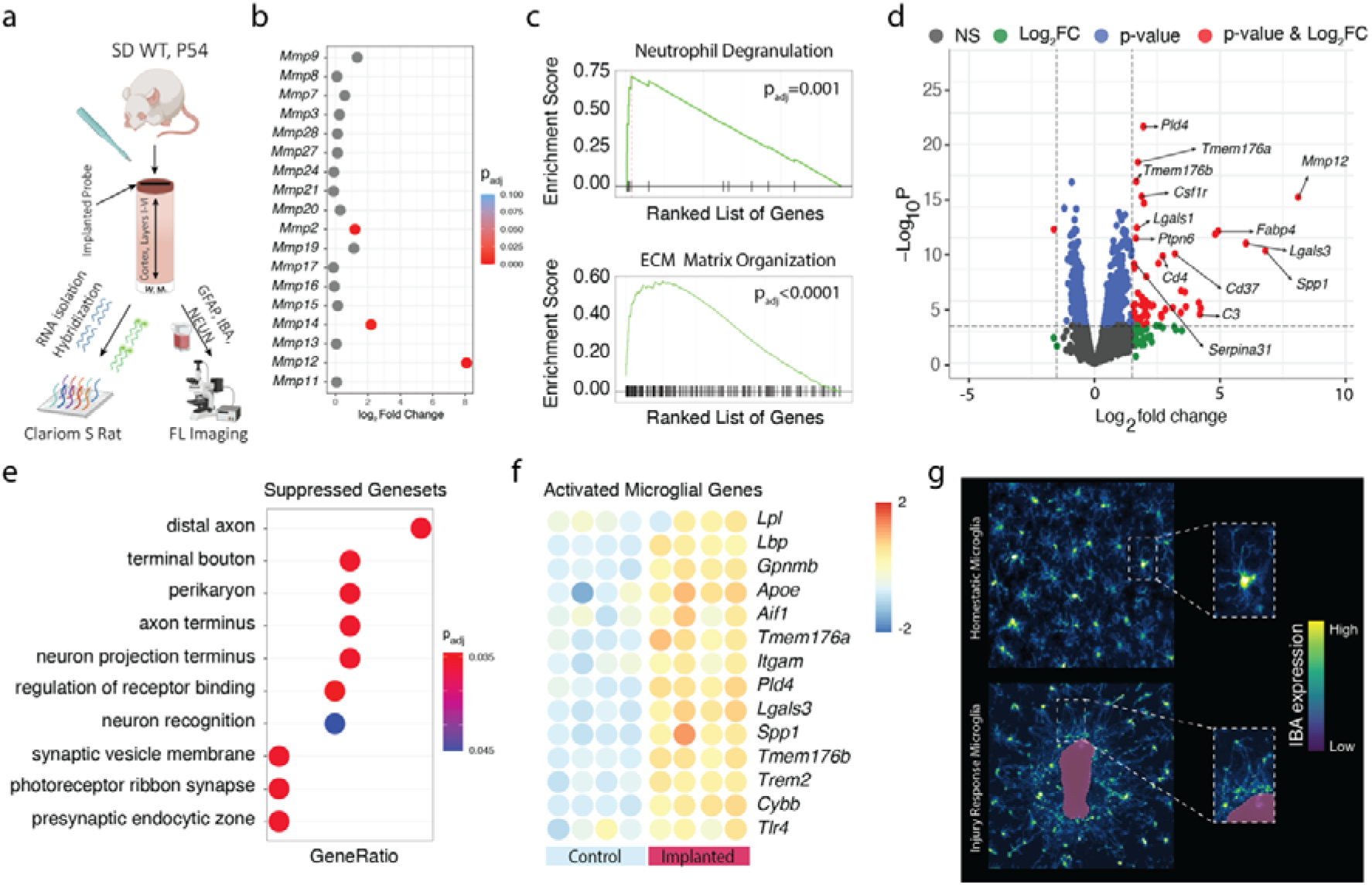
(a) Workflow used for this study (b) Bubble plot of metalloproteinase gene expression 4 hours after probe implantation (c) GSEA results of neutrophil degranulation and ECM matrix organization 4h post implantation (d) Volcano plot of gene expression. Dotted lines represent the cut-off used for analysis (p=0.05, FC=1.3) (e) Dotplot of genesets that are suppressed post implantation (f) Heatmap of reactive microglia genes (g) Immunostaining for Iba^+^ reveals microglia in both homeostatic state and around a sham-implantation site, 4 hours post implantation. Scale bar is 50μm

Probe implantation is associated with significant neural trauma, which is reflected by GSEA results, which shows suppression of gene sets related to neuronal damage, with significantly reduced expression of distal axon, terminal bouton and axon terminus genes, **Fig 1e.** This damage to neurons is accompanied by increased expression of phosphatidylserine (*Ptdss1*, p=0.02), which is a robust indicator of neurons undergoing Wallerian degeneration (45–47). Correspondingly, we report an upregulation of both Apolipoprotein E (*Apoe,* p_adj_=0.002), expressed predominantly by astrocytes, and microglia expressed Triggering receptor expressed on myeloid cells 2 (*Trem2*, p_adj_<0.0001), which have been previously reported to be involved in the sensing of apoptotic neurons, inducing a switch to a neurodegenerative phenotype (48). We also show a significant upregulation of *Stat6* (p_adj_=0.001), which has been reported to orchestrate the clearance of dying neurons in the ischemic brain (49). Overall, we report upregulation of several genes related to the reactive transformation of microglia to a disease associated phenotype of microglia, such as *Lbp*, *Itgam*, *Aif1*, *Tlr4*, *Pld4* and *Tmem176a/b, Lpl, Cybb*, *Gpnmb, Lgals3* and *Spp1*, **Fig 1f** (50). Immunostainings reveal that Iba^+^ (Gene: *Aif1*) cells show a change in cellular morphology, with microglia in the immediate vicinity of the implantation site extending their processes towards the injured region to begin the injury repair process, **Fig. 1g**.

### Reactive astrocyte marker expression within 4 hours of implantation

Reactive astrocytes in the vicinity of a stab wound injury have been reported to appear and persist during the acute phase of implantation (0-2 weeks post cortical breach) (51). Microglial activation, as reported in the previous section, has been reported to be highly correlated with astrocyte activation (52). Significant upregulation of reactive astrocyte markers such as *Gfap*, *Serping1*, *Aspg*, *Fbln5*, *Prelp*, *Phyhd1* and *Tapbp* was observed within 4 hours of implantation, **Fig 2a**. This reactive state is driven by an enrichment of the JAK-STAT pathway (GO:007259, NES=2.04, p_adj_=0.001), which has been previously shown to be involved in the regulation of astrocytic activation (52), **Fig 2b**. In addition, we report a strongly increased expression of complement factor 3, (*C3*, p_adj_=0.003588) that has been shown to be a strong indicator of neurotoxic reactive astrocytes within the active zone of the injury, **Fig. 2a**. This phenotypic switch is reflected in immunostainings, where the highly organized layout characteristic of homeostatic astrocytes is lost, and they start to aggregate around the wound site, **Fig. 2c.**

**Figure 2:**
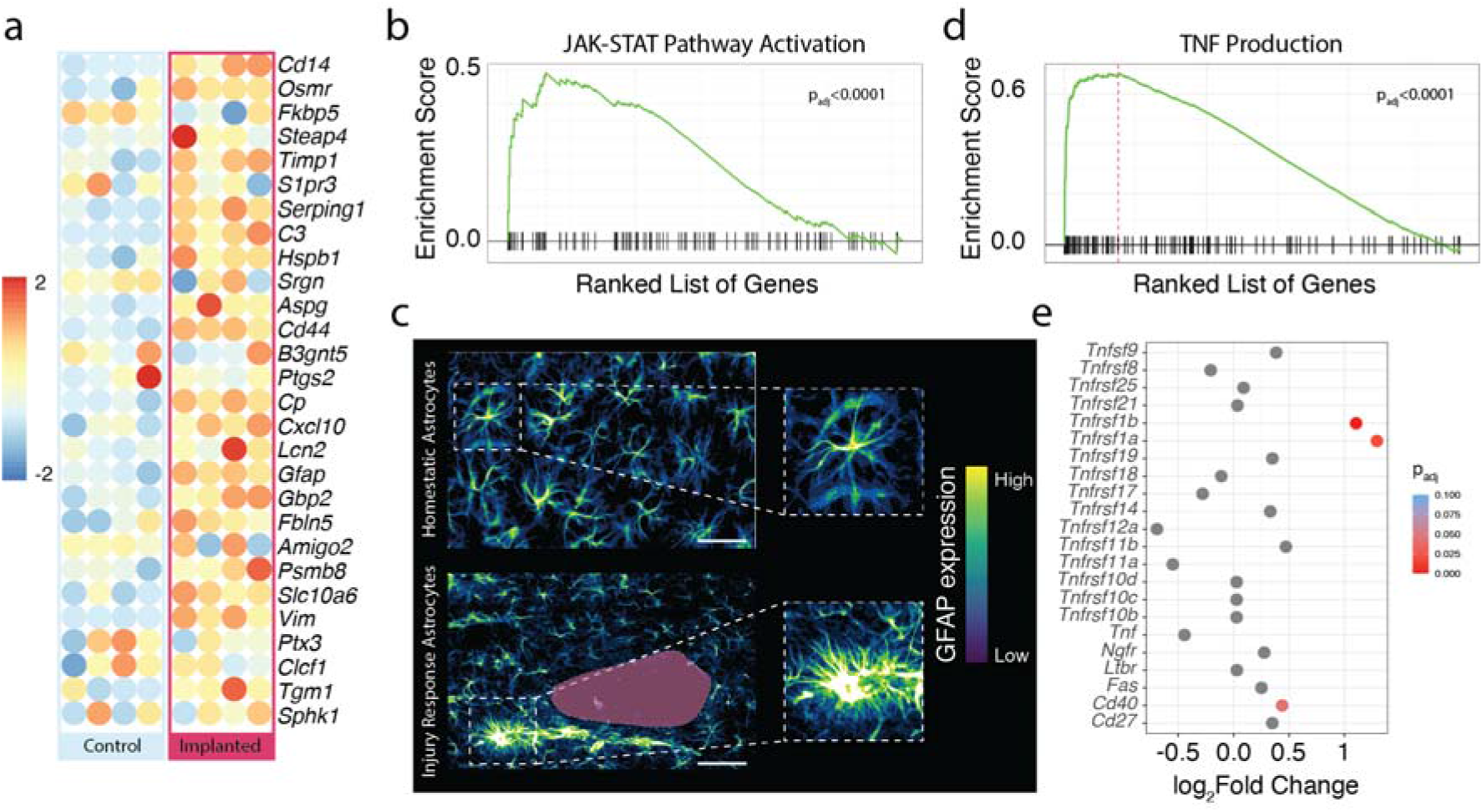
(a) Heatmap showing expression of reactive astrocyte genes (b) GSEA of JAK-STAT pathway activation (c) Comparison between homeostatic and reactive astrocytes in the vicinity of the implantation. Implantation represented in pink (d) GSEA result of TNF Production (e) Bubble plot of Log_2_-fold change of TNF receptor genes. Scale bar is 50μm

The transformation from homeostatic to reactive astrocytes has been shown to be driven by several factors, including Interleukin 1α (*Il1a*), tumor necrosis factor (*Tnf*) or Complement component 1, subcomponent q (*C1qa*). We saw a significant upregulation of *C1qa* (p_adj_<0.0001) and not *Il1a* (p_adj_=0.864) and *Tnfa* (p_adj_=0.717)**, Fig 2e.** *C1qa* has been shown to be dominantly produced by microglia and plays an important role in astrocytic activation (53,54). Even though no significant upregulation of *Tnfa* was observed, a strong enrichment of Tumor necrosis factor (TNF) production (GO:0032640, NES=2.388, p_adj_<0.0001) was seen, and a significant upregulation of a few TNF receptor subtypes was observed (*Tnfrsf1a*, p_adj_=0.02; *Tnfrsf1b,* p_adj_=0.001). Neurotoxic reactive astrocytes are also implied in the loss of synapses and subsequently in decreased frequency and amplitude of miniature excitatory postsynaptic currents, which implies the loss of synaptogenesis(55). Correspondingly, GSEA analysis of our dataset revealed a significant downregulation of genesets related to synapse related processes t, which taken together suggest a loss of synapses due to the implant trauma and subsequent astrocytic activation, **Fig. 1e**.

### Chronic effects within the implantation microenvironment

To study the transcriptomic profile of the brain that is responding to the presence of a foreign object within the neural environment, gene expression profiling was carried out at 1-week, 2-weeks and 4-weeks post implantation. Since the implantation process results in significant damage to the brain parenchyma and to the vasculature, there is an upregulation of genes related to sprouting angiogenesis (GO:002040, NES=1.837, p_adj_=0.001), indicating the start of the repair process of the vasculature. This enrichment is not observed acutely post injury, but only at 1 week post injury. This is coupled with an enrichment of axon guidance genes (GO:0008045, NES=1.96, p_adj_=0.01), which has been previously reported to be strongly correlated with angiogenesis (56). ECM repair was upregulated only for 1 week post implantation, signaling that damage to the neural fabric was quickly repaired by glial cells, to facilitate neuronal repair, **Fig. 3a**. Toll-like receptor signaling, which has been shown to play an important role in the regulation of the inflammatory response in the ischemic brain (57), is significantly enriched at the acute (GO:0002224, NES=2.2468, p_adj_<0.0001) and 1 week (GO: 0002224, NES=1.86, p_adj_=0.006) time point after implantation. A significant suppression was observed 2 weeks post implantation (GO: 0002224, NES=-2.55, p_adj_<0.0001), likely to halt the inflammatory response, which was completely lost at later time points, **Fig. 3b**. This is paralleled by a decline of astrocyte reactivity, which can be observed from immunostainings detailing the increased number of both Gfap^+^ astroglia and Iba^+^ microglia in the vicinity of the implanted electrode, which is reduced at week 2 post implantation, **Fig. 3c-d**. This shows that local regulation of the glial response takes place to limit the harmful effects that reactive glial cells have in the brain.

**Figure 3:**
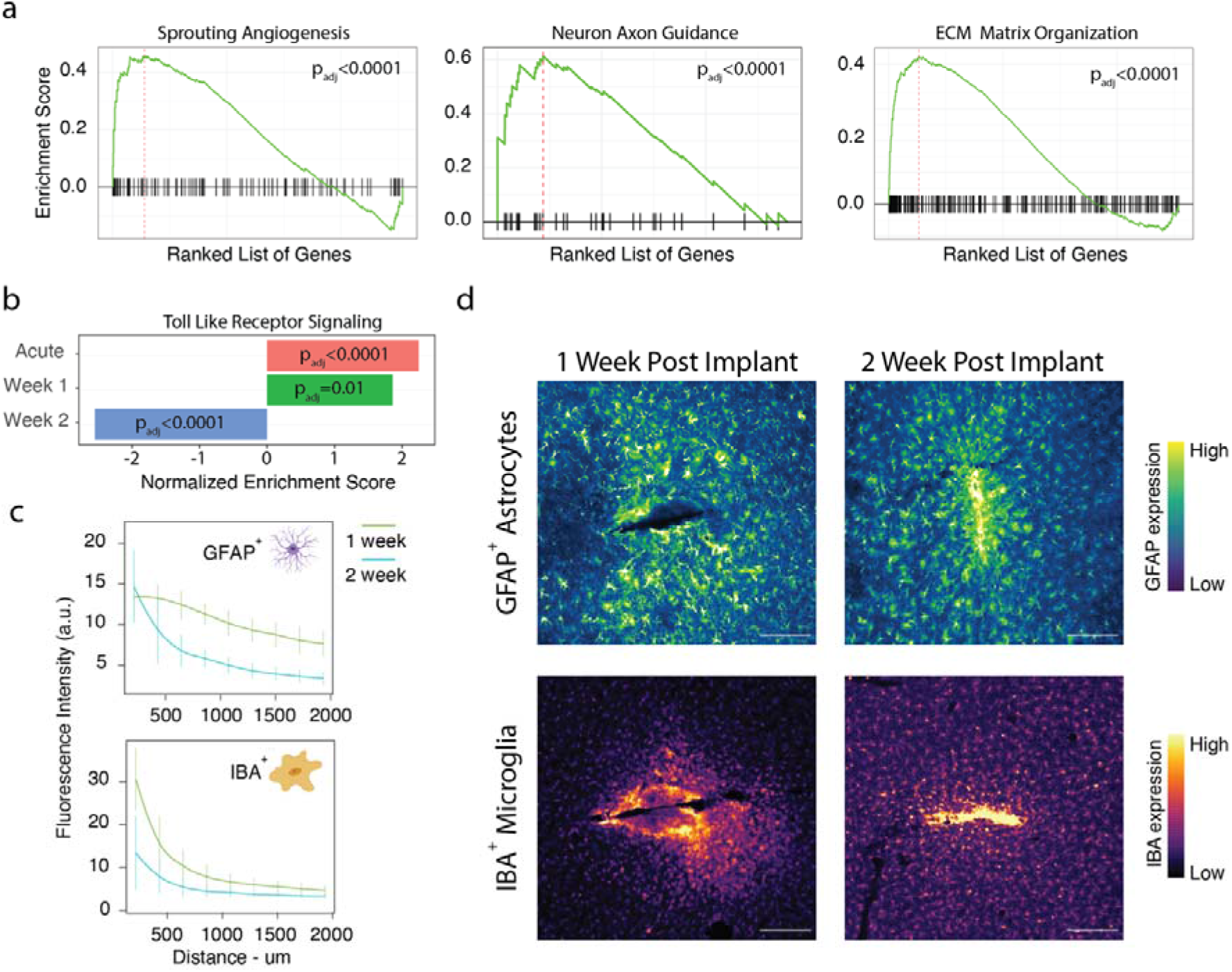
(a) GSEA results showing an enrichment of genesets for Sprouting Angiogenesis, Neuron Axon Guidance and ECM Matrix Organization (b) Change in geneset enrichment for Toll like receptor signaling over 2 weeks post implantation (c) Quantification of the fluorescence intensity for GFAP^+^ astrocytes and IBA^+^ microglia as a function of distance from the implantation site. (d) Representative images of GFAP^+^ astrocytes and Iba^+^ microglia at two timepoints post implantation. Scale bars are 200μm.

Since the presence of activated microglia can be detrimental to the neural environment, therefore the regulation of the microglia phenotype back to the default homeostatic state is essential for recovery (58,59). This change in phenotype is observed 2 weeks post implantation, where our data demonstrates an upregulation of Interleukin 13 (*Il13*, p=0.006), which has been shown to ameliorate neuroinflammation and promotes functional recovery (60). Return of microglia to homeostasis is signaled by the secretion of IL10, mediated by heme-oxygenase^+^ (HMOX1) microglial cells (19). HMOX1^+^ microglia play an important role in the clearance of heme that is released from damaged red blood cells, which plays a role in sterile tissue injury response (61). We report that there is an strong increase of *Hmox1* expression at 4 weeks post injury (p_adj_<0.0001), indicating the return of microglia to the homeostatic state at this time point, which in turn could act to return reactive astrocytes to a homeostatic state as well (19). *Hmox1* is suggested to aid in resolving neuroinflammation and has been found to be necessary to attenuate neuronal cell death, and the clearance of blood from the parenchyma (61). Complementing this return to homeostasis, we report increased expression of *Socs3*, which further aids in the downregulation of inflammation in both macrophages and microglia (62). Neuronal repair mechanisms are not complete at this time point, and correspondingly, significant upregulation of *Tyrobp* (p_adj_=0.0001) is still seen at 4 weeks post injury, reported to be expressed by microglia involved in injury repair (63), **Supp. Fig. 1a**. This is coupled with a significant enrichment of genes expressed by neuroprotective astrocytes, including *S100a10*, *Emp1*, *Mobp* and *Cd14*, *Gfap*, *Timp1*, *Hspb1*, *Cd44*, *Cp* and *Vim* were significantly upregulated at this time point, **Fig. 4a**. This is indeed the case, where we see a significant enrichment of *Gfap*^+^ astrocytes around the implant scar, highly overlapping with NeuN^+^ neurons, **Fig. 4b.**

**Figure 4:**
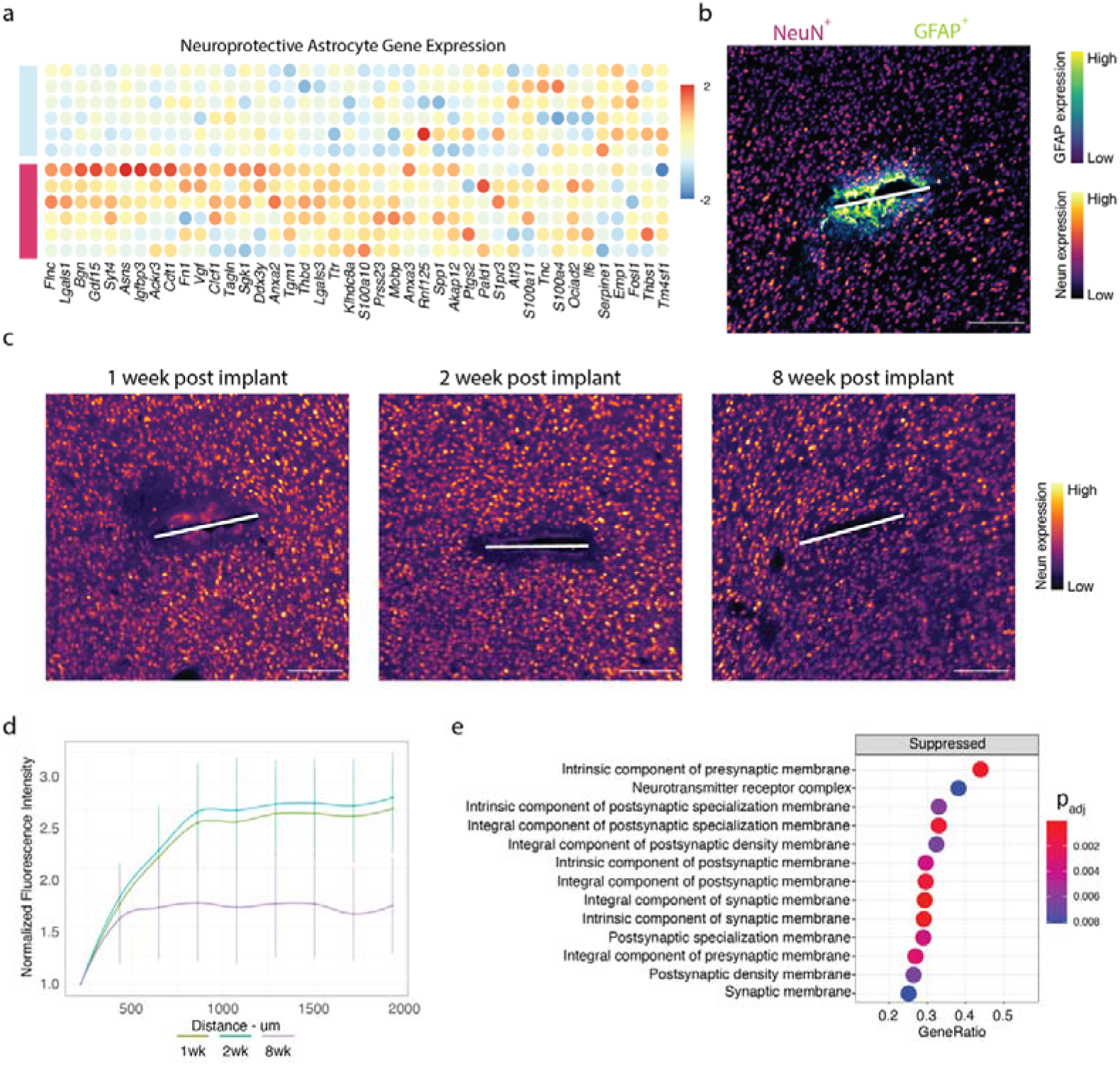
(a) Heatmap of neuroprotective astrocyte gene expression at 4 week time point (b) Coregistration of NeuN^+^ neuronal cells and GFAP^+^ astrocytic cells at the implant site at 4 weeks post implantation (c) Representative images of NeuN^+^ immunostainings show neuronal cell recovery at the electrode implantation site over 8 weeks of implantation (d) Quantification of NeuN^+^ fluorescence intensity from the implantation site at different time points (e) Dotplot of suppressed genesets at 4 weeks post implantation, obtained using GSEA.

These neuroprotective astrocytes play an important role in the remyelination of damaged axons, by secretion of Interleukin 1B (*Il1b*, p_adj_=0.0006) and tissue inhibitors of metalloproteinases, (*Timp1,* p_adj_<0.0001), implicated in the proliferation and differentiation of myelinating oligodendrocyte cells (64). We also report an upregulation of Thrombospondin1 (*Thbs1*, p_adj_=0.004), which has been shown to be involved in the formation of excitatory neurons. This further indicates the role of the neuroprotective astrocytes, inducing repair mechanisms for damaged neurons (65), **Supp. Fig. 1a**. This is also reflected in immunostainings showing that compared to earlier time points, NeuN^*+*^ cells are significantly closer to the implantation site 4 weeks post implantation, **Fig. 4c-d**. Correspondingly, a significant loss of genes related to synaptic compartments was seen, hinting at a loss of synaptic features, **Fig. 4e.**

### Transcriptomic profile of chronic implantation

To understand the long-term effects of chronic cortical electrode implantation in the neural environment, a final cohort of 6 rats were left implanted for 18 weeks post-surgical intervention. Although a strong upregulation of inflammatory pathways was observed throughout the time span presented in this work, we wanted to determine the time scale at which the transcriptional profile would resemble non-implanted conditions. Current understanding assumes the brain to return to a homeostatic state within 12 weeks of electrode implantation (66). However, in our work, we present evidence that the wound healing process is not complete (GO:0042060, NES=1.6, p_adj_=0.003) at 18 weeks post implantation, even though we made use of a probe with low bending stiffness. Additionally, complement activation remains significantly enriched even at this time point, (GO:0030449, NES=1.6, p_adj_=0.001). The presence of the probe over chronic time periods is in agreement with a recent review pointing to chronic activation of complement pathways in other chronic neurodegenerative disorders, such as MS and AD (67–70), **Fig. 5a**. The significant increase in expression of metalloproteinases that was seen at acute time points was lost at 18 weeks post implantation, signifying that the ECM degradation is under control, as expected at this late time point, **Supp. Fig. 1b**.

**Figure 5:**
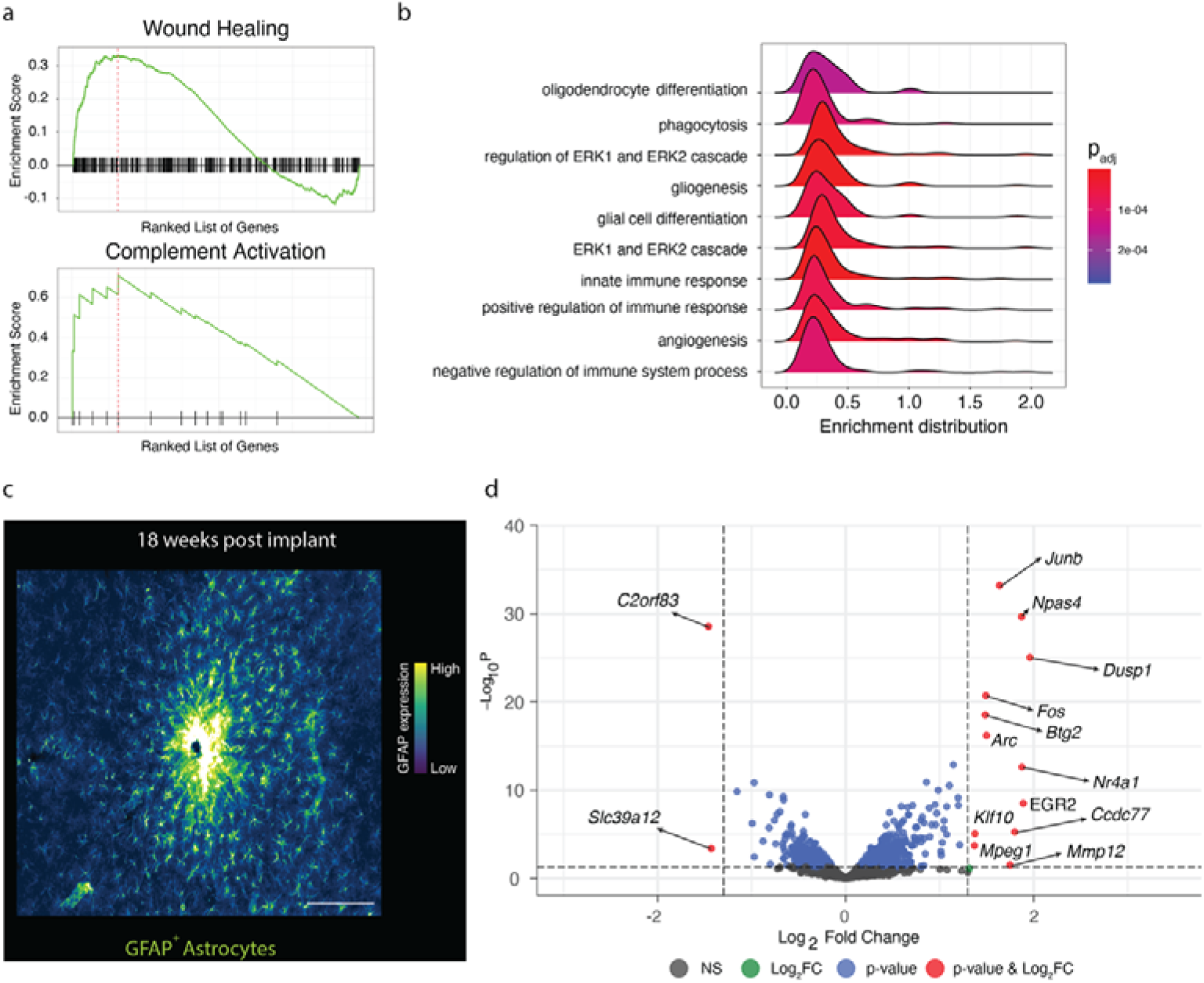
(a) GSEA results showing upregulation of wound healing and complement activation pathways at 18 weeks post implantation (b) Ridgeplot showing enrichment of genesets significantly upregulated at 18 weeks post implantation (c) Representative image showing the astrocytic scar present at 18 weeks post implantation. Scale bar is 200μm (d) Volcano plot showing significantly up- and downregulated genes at 18 weeks post implantation. The dotted lines represent cut offs used to detect significant differentially expressed genes.

Btg2 is highly expressed (padj<0.0001) at this time point, pointing to ongoing neurogenesis, identifying neuroepithelial cells that switch to neuron generating division(71). One observation of interest we report is the significant enrichment of oligodendrocyte differentiation, gliogenesis (GO:0042063, NES=1.97, p_adj_<0.0001), and ERK1 and ERK2 cascade (GO:0070371, NES=1.93, p_adj_=0.0001), which could be indicative of persisting astroglial activation, ongoing re-myelination and release of growth factors, **Fig. 5b**. This is indeed the case, with increased numbers of GFAP^+^ astrocytes still accumulating around the implantation site, **Fig. 5c**. Most strongly upregulated genes at this time point are the immediate early genes *Junb*, *Fos*, *Btg2*, and *Arc* (Activity Regulated Cytoskeleton Associated Protein) which may reflect hyperactivity of neurons in the vicinity of the implant. Interestingly, *Npas4* (Neuronal PAS Domain Protein 4) is also significantly upregulated (padj<0.0001). This transcription factor is expressed by neurons and has been implicated in regulating the excitatory-inhibitory balance within neural circuits plays a key role in the structural and functional plasticity of neurons.

## Discussion

In the context of implantation of electrodes into the brain the glial scar has been a major topic of interest in the past since the effectivity of brain machine interfaces is critically depends on the quality of the neuronal activity recorded which is inversely correlated to the extent of glial scaring. Most studies to date have relied on immunocytochemistry with antibodies against the major cell types of the nervous tissue to assess the response of neurons, astrocytes and microglial cells to the foreign body. More recently broader analyses have been performed with qPCR, focusing on involved neuroinflammatory pathways known to be involved in regulating glial scarring (72) and RNA-sequencing to determine expression changes of a larger set of genes around an implanted probe. To capture the full complexity of the response of the tissues upon probe implantation as well as the underlying molecular mechanisms, we carried out unbiased transcriptomic analyses over an extended period of time after implantation of a flexible electrode. We see that the brain employs the same mechanisms for glial scar formation, as has been previously reported. Within 4 hours of introducing a foreign body being into the brain, vascular disruption leads to local bleeding with extravasation of blood components and blood cells which triggers activation of microglia. This microglial activation in turn induces astrocyte activation. Genes which have been reported to be indicative of the inflammatory, neurotoxic A1 phenotype of astrocytes (43) such as *C3*, *Serping1*, *Gbp2*, *Fbln5*, *Psmb8* are robustly upregulated within 4 hours after probe implantation. Similar to what has been reported by Thompson et al., 2021, components of the TNF-signaling pathway, e.g., Tnfrsf1a and b are up-regulated at this early time point. Tnfrsf1a has been shown to activate the NF-κB and MAPK-pathway in microglial cells, known components of inflammatory signaling (62). Expression of complement C3 and C1q is also upregulated already at the 4h post-implantation time point and have been suggested to play a role in neural synapse elimination in pathological conditions (63). Over time, this inflammatory reactive transformation of astrocytes is lost, and is replaced by neuroprotective, scar forming astrocytes within one week of implantation and maintained over the whole study period (18 weeks). Together with reactive microglia these neuroprotective astrocytes are critical for neuronal repair. Microglia present in the vicinity of the implantation regulate the repair to the ECM and the reactive states of astrocytes, mediated by release of IL10 cytokine. The infiltrating macrophage/microglia population plays a role in restoring the homeostatic state by reactive polarization to enable tissue repair. Our data shows that there is an increase in *Hmox1^+^* microglial cells at 4 weeks post implantation, signaling an end to the inflammatory environment and ushering an anti-inflammatory state. This state persists till the end point in this study, with persistent gliosis occurring till 18 weeks post implantation. From a biological standpoint, this is one of most important findings we present. Even after 18 weeks of implantation, we saw significant enrichment of genes related to gliogenesis and glial cell differentiation, despite flexible brain implants, confirmed by immuno-fluorescence imaging. Based on the Cancer mine atlas, an expert curated database of oncogenes, we see that most of the genes upregulated at this timepoint were known oncogenes and could indicate an alarming development, involving cortical implants being a “seed” for further disorders of glial development, as seen with other chronic neurodevelopmental conditions such as MS and AD. However, the timeline that we have presented in this work is limited to 18 weeks and making concrete claims about the potential oncogenic consequences of cortical implants would be difficult to justify. On a final note, this work helps us to map the transcriptome in terms of potential cellular interactions possibly to be interrupted to modulate the immune responses that occur due to bioelectronic interfaces ensuring optimal signal fidelity over chronic timescales. Taken together our study, to the best of our knowledge, is the most comprehensive in both depth and observational time span, describing molecular and morphological changes of nervous tissue to implantation and presence of thin, flexible intraneural probes. Knowledge of key molecular mechanisms and signal transduction pathways associated with intraneural probes will help to design identify the most relevant responses which leads to probe failure in the long run. Based on the knowledge of the most strongly affected genes and pathways, a panel of biomarkers can also be used to help develop more compliant probes in the future.

## Acknowledgments

Elements from Figure 1 and Figure 3 were created using Biorender.com. Part of this study was supported by the German Ministry of Research project FMT 13GW0230A and the German Research Foundation’s (DFG) Cluster of Excellence BrainLinks-BrainTools (EXC 1086).

## Author Contributions

KJ, MJ, CM and MK carried out experiments and data acquisition. KJ carried out RNA and imaging data analysis. MK, ST, TS, CAH and UGH conceived and supervised the project and provided resources. KJ and UGH wrote the initial draft. All authors discussed the results and were involved the preparation of the final submission draft.

## Data Availability

The raw data required to reproduce these findings cannot be shared at this time as the data also forms part of an ongoing study. The processed data to reproduce the findings are part of the supplementary materials.

## Notes

DISCLOSURE OF CONFLICTS OF INTEREST: No potential conflicts of interest are disclosed by the authors.

### Competing Interest Statement

The authors have declared no competing interest.

